# Spawning asynchrony and mixed reproductive strategies in a common mass spawning coral

**DOI:** 10.1101/2024.12.06.627154

**Authors:** Gerard F. Ricardo, Christopher Doropoulos, Russell C. Babcock, Arne A. S. Adam, Elizabeth Buccheri, Natalia Robledo, Julian Uribe-Palomino, Peter J. Mumby

**Affiliations:** Marine Spatial Ecology Lab, School of the Environment, The University of Queensland, St. Lucia, QLD, Australia; CSIRO Environment, St. Lucia, QLD, Australia

## Abstract

Understanding strategies of organisms that utilise multiple modes of reproduction presents a complex challenge for evolutionary biologists. Platygyra daedalea, a common reef-building coral with unclear reproductive boundaries between morphological species, illustrates these complexities. Here, we evaluate the contribution of these reproductive modes in the coral P. daedalea at Heron Island, on the southern Great Barrier Reef. We tagged and sequenced eighteen coral colonies, representing various degrees of spatial clustering, along a 130-m stretch of shallow reef slope. During spawning, divers collected eggs from each colony and placed them in mesh containers within the spawn slick, allowing free movement of sperm for fertilisation. High levels of spawning asynchrony were observed, potentially indicating distinct genetic clusters within the putative species, which resulted in low fertilisation success (1.5%). Notably, of those fertilised eggs, paternity assignments revealed that all resulting embryos were self-fertilised, with no cross-fertilisation occurring. The adult population showed evidence of two genetically distinct subpopulations, along with levels of spatial autocorrelation and inbreeding. This evidence supports the notion of small breeding populations within larger assemblages, density-dependent population effects, and localised recruitment. Selfing may serve as a reproductive assurance mechanism in such populations, which may be more hierarchically structured than previously thought. Given the lack of evidence for in situ outcrossed fertilisation in this natural coral population during split spawning, it appears that P. daedalea may rely on limited high-density patches of adults for successful cross-fertilisation, utilise atypical modes of reproduction when at low densities, and/or sustain its population through limited progeny.

## Introduction

Colonial organisms often employ multiple reproductive strategies to sustain their population in dynamic and competitive environments. Benthic colonial invertebrates share many similarities with higher plants, such as a modular construction, sessile adult stages, hermaphroditism, synchronous external reproductive events that produce masses of progeny (termed ‘mast seeding’ in plants (Kelly 1994) and ‘mass spawning’ in benthic invertebrates (Harrison et al. 1984)). Sessile organisms face unique challenges related to mating including how to achieve fertilisation and exercise mate-choice or optimise sex allocation (Serrão & Havenhand 2009). A central theory explaining the persistence of colonial organisms is through the use of multiple modes of reproduction under evolutionary and generational time scales or in response to different environmental conditions (Williams 1975, Barrett 2014, Uecker 2017). Outcrossed sexual reproduction often enhances dispersal and genetic diversity leading to greater tolerance to disturbances at the expense of higher energetic demands through meiosis and gametic waste.

In broadcast-spawning invertebrates, increased spacing between individuals and the dilution of gametes can reduce the likelihood of gamete encounters, leading to lower reproductive success, a component Allee effect (Levitan & Petersen 1995, Babcock & Keesing 1999, Courchamp et al. 1999, Knowlton 2001). Similarly, at least 10% of angiosperms rely on wind pollination, a process that carries a similar risk of low fertilisation from pollen limitation (Friedman & Barrett 2009). In wind pollinated plants, critical distances for reduced fertilisation can exceed 60 m (Knapp et al. 2001), whereas in sessile marine invertebrates, this threshold is poorly understood and could be restricted to just tens of metres (Coma & Lasker 1997, Teo & Todd 2018, Mumby et al. 2024). In contrast to terrestrial systems, our understanding of reproductive isolation and the dynamics of mixed reproductive modes in marine invertebrates remains limited, highlighting a critical gap in our knowledge of coral reproductive ecology.

Self-fertilisation, or selfing — a mode of sexual reproduction between a single individual’s gametes — is common in simultaneous hermaphroditic organisms (Jarne & Auld 2006). Approximately 15– 20% of hermaphroditic animals are highly selfing (Jarne & Auld 2006), as are 10–15% of flowing plants (Wright et al. 2013). Selfing provides reproductive assurance in the absence of mates, helping to maintain adaptive genes suited to local conditions. Over evolutionary timescales, shifts from outcrossed to selfing-dominated reproductive modes are common. However, selfing lineages tend to be short-lived as they eventually face challenges associated with low genetic diversity, such as inbreeding depression and reduced capacity to respond to environmental disturbances (Goodwillie & Weber 2018). Likewise, asexual reproduction typically allows rapid expansion at smaller spatial scales, but the propagation of successful genotypes at the expense of diversity leads to greater extirpation risk during environmental stress (but see Kirkpatrick and Jarne (2000)). Species that rely on asexual modes, typical of colonising species, are subject to frequent reductions in population densities (Manning 1981), but nevertheless can be successful. In some cases, asexual clones can reach staggering sizes: a polyploid clone of seagrass in Western Australia has reached 180 km and survived for 8500 years (Edgeloe et al. 2022).

Different modes of reproduction generate varying levels of genetic diversity through distinct reproductive processes. Outcrossed sexual reproduction increases genetic diversity through the process of meiosis and the fusion of chromosomes from different parents. Selfing, similarly allows for some genetic diversity through the processes of recombination and independent assortment of chromosomes during meiosis, but is more limited, as the fusion involves chromosomes from the same parent (Hartl 1997). While selfing rate estimates are often inferred from indirect population-level metrics, these metrics can be heavily biased if assumptions about population structure, inbreeding and presence of genotyping errors are not met (Bürkli et al. 2017). Data from progeny arrays using genetic markers provide a direct estimate of selfing with fewer assumptions compared to population-level metrics. However, progeny arrays are rarely conducted in animals, especially in situ, because of their cost- and labour-intensive nature (Hartl 1997, Coffroth & Lasker 1998, Jarne & Auld 2006). Asexual reproduction, such as fragmentation, typically results in progeny with the same genomic arrangement as the parent, apart from rare mutations. Parthenogenesis, a type of asexual reproduction where an egg develops into an embryo without sperm entry, is widespread in terrestrial taxa such as plants, insects and reptiles (Bierzychudek 1985, Watts et al. 2006, Normark & Kirkendall 2009), but poorly understood in marine invertebrates. Parthenogenesis can result via mitosis or meiosis leading to an absence or small amount of genetic variation.

While much of the work on the evolutionary roles of mixed reproductive modes has been conducted in higher plants, sessile marine invertebrates such as corals offer alternative and underutilised model organisms to examine these roles. This is due to their diverse reproductive strategies, sessile nature, and general interest how they may adapt under broad scale disturbances such as climate change. Approximately 80% of corals release gametes into the water column (Harrison 2011), allowing for random mating and dispersal. Brooding, which occurs in about 17% of coral species and can involve either sexual or asexual reproduction, is a less common reproductive strategy. While selfing is generally considered uncommon in corals, there are a few notable genera that exhibit high levels of selfing such as Goniastrea, Favia, Seriatopora, Platygyra, and Porites (Willis et al. 1997, Gleason et al. 2001, Miller & Mundy 2005). Various modes of asexual reproduction that result in clonal propagation exist in corals such as fragmentation of various life history stages (Wallace 1985, Babcock 1991, Heyward & Negri 2012), polyp-bailout (Sammarco 1982), and parthenogenesis (Combosch & Vollmer 2013, Vollmer 2018).

Previous studies have documented a range of in situ fertilisation outcomes influenced by spatial population metrics, species traits and hydrodynamic factors (Coma & Lasker 1997, Levitan et al. 2004, Miller & Mundy 2005, Mumby et al. 2024, Ricardo et al. 2024). However, it remains unknown whether successful fertilisation can occur during split spawning events, where altered gravid population densities may influence gamete concentrations and fertilisation success. Split spawning— where individuals at a given location release gametes across multiple months—provides a unique natural experiment to examine shifts in reproductive strategies under variable environmental conditions. Although split spawning has been hypothesised to realign reproduction with favourable conditions (Willis et al. 1985, Foster et al. 2018), it may also elevate the risk of fertilisation failure, particularly in low-density populations, by further reducing the effective pool of gametes. In this study, we use the coral Platygyra daedalea as a model to investigate mixed modes of reproduction during a split spawning event. While selfing occurs in Platygyra at elevated levels relative to other coral genera, outcrossing remains the primary mode of reproduction (Willis et al. 1997). To investigate these reproductive modes, we employ next-generation sequencing for paternity assignments and population structure analysis to directly assess the relative contributions of selfing, outcrossing, and asexual reproduction.

## Methods

### Coral and site selection

The genus Platygyra is known as a syngameon, with unclear reproductive boundaries between morphological species (Miller & Babcock 1997). Platygyra daedalea is primarily a hermaphroditic submassive coral that usually spawns six to eight days after the full moon on the southern GBR (Baird et al. 2021). Egg-sperm bundle setting can be difficult to observe, and the species is known to have spawning times ranging from dusk until several hours after sunset (Miller & Babcock 1997). The colony releases egg-sperm bundles instantaneously, and egg-sperm bundles contain ∼60 eggs per bundle, with eggs being ∼400 µm in diameter (Babcock et al. 2003).

Corals of P. daedalea were examined for presence of pigmented eggs starting two days prior to the full moon on the 8^th^ of December 2022. As corals on the mid-shelf and outer Great Barrier Reef typically spawn in November, a split spawning was predicted owing to the early full moon occurring in December, increasing the risk of Allee effects occurring during fertilisation. A preliminary survey revealed a very low proportion of gravid corals in most genera, with approximately 25% of the P. daedalea population on the reef slope showing mature oocytes. Our chosen field site was situated at Coral Canyons (23.4548° S, 151.9238° E), located on the western side of Heron Reef, adjacent to the Heron-Wistari channel. This location is marked by distinctive tidal currents along the reef slope, flowing northwest during flood tides and southeast during ebb tides. The site is exposed to waves during common south-easterly swells, median peak period of 4.7 s and a median significant wave height of 0.76 m (Fig. S1). All corals along 150 × 36 m of the reef slope were assessed for gravidity by progressively shearing small layers of tissue off the coral’s surface to expose the egg layer. Only gravid colonies from the study site with a mean diameter of ≥ 10 cm were included, and a total of 19 gravid corals were tagged along the reef slope, which extended from ∼2 m to 9 m depth, with 15 monitored during spawning periods. Most morphotypes were identified as P. daedalea PDC (classic morph) following descriptions by Miller (1994a). Platygyra species are known to create hybrid embryos, but only one other species was found in the patch (Platygyra sinensis) that was not gravid. During daylight hours, each tagged colony was georeferenced using a handheld GPS (Garmin eTrex 10). Colony size was measured by taking an image of the colony from directly above and using a graduated diving rod as a scale. A marking line was run sequentially between each gravid colony, enabling divers to quickly identify any indications of spawning (see below).

### Experimental design

To assess in situ fertilisation success across the transect, we constructed spawn collectors by mounting a funnel above the coral using metal wire frame. The funnel channelled gamete bundles into a detachable Falcon tube (herein ‘mesh containers’). A portion of the container was cut open and covered with 150 µm mesh to allow for sperm to move freely through the containers while retaining the individual colony’s eggs. Glowsticks were securely fastened to the mesh containers for easy tracking during the night. Movement of sperm through the mesh to fertilise eggs has been validated by Ricardo et al. (2024), confirming the effectiveness of this approach.

During the predicted coral spawning window, divers were deployed at sunset (∼18:00) and corals monitored along the marking line using red-beam dive torches. When spawning was observed in individual colonies, the mesh containers were removed from the collector, capped, and their attached glowsticks were activated before being released. Released mesh containers were tracked for over 80 minutes, with their collection locations marked using GPS. After retrieval, the mesh containers were transferred to sperm-free seawater collected prior to spawning and transported to Heron Island Research Station for assessment. Gametes within each mesh container at ∼3 hours post spawning were scored as a proportion of the total unfertilised eggs and embryos. These embryos were then isolated into individual wells of 6-well plates containing filtered seawater for development into larvae for later genetic work. Embryos were isolated at ∼4-cell stage, before they become fragile, thereby preventing the risk of fragmentation (Ricardo et al. 2016).

In addition to the main experiment, other modes of reproduction were assessed ex situ to assess self-fertilisation and parthenogenesis. To assess self-fertilisation, two colonies outside of the patch were collected and brought back to aquaria facilities at Heron Island Research Station. A single colony spawned on the 14^th^ of December 2022 and used as a self-fertilisation control. The eggs were left in ∼10^7^ sperm mL^-1^ and assessed at three- and six-hours following spawning. During the following year, parthenogenesis was assessed by collecting egg-sperm bundles from two isolated colonies of P. daedalea. Bundles were separated into eggs and sperm using a 100-µm mesh. Eggs were washed with filtered seawater and divided into containers (n = 3 per individual) containing 100 mL filtered seawater. The eggs were assessed the following day for indications of fertilisation via observations of cell division.

### Genetics sampling and analyses

Genetic sequencing was used for paternity assignments and to assess population structure. A small amount of tissue from each of the monitored 15 colonies, and an additional three unmonitored colonies, was collected into zip-lock bags for sequencing. Replicate samples (n = 2) of adult coral colony tissue were stored in 100% ethanol at -80°C until further processing in the lab. Embryos underwent development for more than 36 hours before being stored in 10 µL seawater in 100% ethanol solution. The samples were transported to the University of Queensland and stored at -20°C. Prior to sequencing, samples of adults and larvae were transferred to 96-well plates, capped, and sent to Diversity Arrays Technology Pty Ltd (Canberra, Australia) for DNA extraction, library preparation, and single nucleotide polymorphism (SNP) sequencing. Codominant, genome-wide, biallelic SNP markers were detected at 0.8 mln reads. DArTSeq sequencing is a technique that employs pairs of restriction enzymes to digest genomic DNA, enabling the identification of SNP markers. Adapters are added to the sequence fragments and sequenced on an Illumina short-read platforms. These terminal sequences include a barcode for disaggregating the sequences of each sample during subsequent analysis. The selected DNA fragments are sequenced in a shortened process to produce raw “sequence tags” of approximately 75 base pairs. These tags are then filtered based on sequence quality, especially in the barcode region, truncated to 69 base pairs, and grouped by sequence similarity. A series of proprietary filters are applied to select the sequence tags that contain a reliable SNP marker. One-third of the samples are processed twice as technical replicates, starting from DNA and using independent adaptors, through to allelic calls. High-quality markers with low error rates are primarily selected using scoring consistency (repeatability). Sequencing was conducted against a reference P. daedalea genome v1.0 (Voolstra et al. 2015, Liew et al. 2020).

Raw SNP sequencing data were subjected to standard filtering processes. Loci sharing SNPs were identified and duplicates removed, retaining only one fragment per set. Next, loci were filtered to a minimum read depth of 10. Loci with limited reproducibility were also excluded using a threshold of 95%. Additionally, loci and individuals with low call rates, indicating high proportions of missing data, were filtered out using thresholds of 70% and 58% respectively. SNP markers were filtered by retaining loci where the SNP and reference allele were between 0.01 and 0.99, and where sequencing coverage exceeded 4 reads to ensure the inclusion of polymorphic loci with sufficient depth.

### Paternity assignments and progeny arrays

Paternity assignments were conducted on offspring found within each mesh container. Colonies that contributed eggs and were tracked within mesh containers were hereafter referred to as ‘egg donors’, whereas initially unidentified colonies that provided sperm for fertilisation are hereafter referred to as ‘sperm donors’. As a proof of concept, we tested if paternity assignments of larvae (n = 3) could be determined correctly from a pool of candidate parents (n = 11). Briefly, 11 Acropora kenti (previously A. tenuis) adult colonies were collected from Orpheus (18.6194°S, 146.4859°E) and Pelorus Island (18.5374°S, 146.4934°E) reefs in Nov 2022. Following spawning, a larval culture was created between two known parents. Samples of each colony and three 42-hour-old larvae from the culture were placed in 100% ethanol solution, and stored at -20°C. These samples were sent for SNP sequencing (0.25 mln reads) and basic filtering at DarT P/L as described above. SNP data underwent further filtering that reduced the total number of usable loci to 257, which is typically sufficient to predict parent-offspring pairs in SNP data (Anderson & Garza 2006).

Paternity assignments were carried out using the CERVUS 3.0.7 software (Kalinowski et al. 2007). CERVUS is a simulation-based approach that uses likelihood ratios to assign parentage. The simulation-based approach is relatively robust to null alleles and genotyping errors (Kalinowski et al. 2007). Following assignment of one of the known parents as the dam (egg donor colonies), the following settings were used: simulated offspring = 20,000, candidate sires (sperm donors) = 10, proportion of colonies sampled = 0.99, proportion of loci typed = 0.99, proportion of loci mistyped = 0.01, minimum typed loci = 128. All larvae were assigned correctly to their known sperm donor as the most likely candidate parent (Fig. S2). Although none of the assignments indicated high levels of confidence—likely due to the low-density sequencing used for this preliminary analysis—our tests successfully assigned a candidate sperm donor.

Next, paternity assignments were conducted on P. daedalea larval samples with known maternal genotypes at time of egg capturing. The CERVUS analysis was under strict (95%) and relaxed (80%) confidence levels for assignments. Simulations were based on allele frequencies for the mapped population, involving 10,000 offspring, 80% sampled candidate sperm donor, and a 1% error rate. An 80% sampled candidate sperm donor’s rate was selected because of a small proportion of unsampled colonies that are found on the reef flat upstream of the site. CERVUS cannot identify unsampled candidate parents, but the uncertainty in the simulations is reflected in the level of confidence of the assignments. To increase robustness and certainty, the paternity assignment was repeated using the software COLONY 2.0 (Jones & Wang 2010), using the following inputs: Mating System I = polygamy, Mating System II = inbreeding, clone, Species = monoecious, Analysis Method = Full-Likelihood, with the probability of sperm donors within the patch having equal probability. Further, changes in heterozygosity between known parents and offspring would additionally support selfing (Ayre & Miller 2006), and were compared using *F*_*IS*_ = 1 - *H*_*o*_/*H*_*e*_, where *H*_*o*_ is the observed heterozygosity and *H*_*e*_ is the expected heterozygosity. *F*_*IS*_ between parents and offspring were compared using a t-test after checking for normality and homogeneity of variances. Genetic variation of the larvae in respect to the adults were visualised using Principal Component Analysis (PCA) with the package ade4 (Dray & Dufour 2007).

### Subpopulation structure

Null alleles can falsely increase homozygosity (David et al. 2007), and an additional filtering step was added using the package popgenreport (Adamack & Gruber 2014) for analyses more sensitive to these artefacts. To assess the genetic structure across the adult P. daedalea population at our site, we performed Principal Component Analysis (PCA) ordination combined density-based spatial clustering DBSCAN (Hahsler et al. 2019), or model-based clustering methods using STRUCTURE (Pritchard et al. 2000). For the PCA, the first two principal components were used to assess genetic groups. Appropriate eps values for DBSCAN clustering were determined using k-nearest neighbour distance plots, with the elbow method to identify the optimal threshold. For STRUCTURE analysis, the Evanno method was used to determine the optimal K value for genetic groups using the admixture model. For both analyses, only the highest quality replicate of each unique individual was analysed to prevent artificial clustering. Similarly, the inbreeding coefficient (F), defined as the probability for an individual to inherit two identical alleles from a single ancestor, was calculated with the package adegenet (Jombart & Ahmed 2011).

Linkage disequilibrium was assessed using the Index of Association (IA) using the package poppr (Agapow & Burt 2001, Kamvar et al. 2014). IA indicates the non-random association of alleles between loci. During sexual fertilisation, meiosis shuffles alleles through recombination, leading to random associations of alleles. Therefore, non-random association can indicate the absence of recombination, as seen in mitotic mechanisms such as clonality, but may also indicate subpopulation structure.

### Statistical

Georeferenced individuals were analysed for nearest-neighbour distances and Clarke-Evans index using the package spatstat (Baddeley & Turner 2005) in R (version 4.2.1). A Mantel test was conducted to assess pairwise differences in spawning times between individuals and their pairwise distances using the package vegan (Dixon 2003). A Fisher exact test was used to compare genetic groups with spawning times. Spawning time was binned into categories ‘early’ and ‘late’ using a 45- minute time threshold because exact spawning times were not available for corals that spawned after the divers had exited the water. A t-test was used to compare genetic groups with depth.

## Results

### Environmental conditions

During the two primary spawning days that the experiment was conducted, winds from the north to northwest at ∼20 km h^-1^ (∼11 knots) were typical for the region during spawning (Barneche et al. 2021) (Fig. 1). Tides occurred during the low or early flood stages at the time of spawning, transitioning to a flood tide during the periods of mesh container release (Table 1). Although low to moderate winds prevailed from the N-NW direction, the movement of the slick and containers was primarily driven by tidal currents in a W-NW direction (Fig S2b). The containers moved at a speed of 0.05 m s^-1^ on both nights (approximately 270 m during the 90-minute release). On the second night of spawning, the final positions of the containers were aligned parallel to the reef at 161.2 m ± 10.3 from shore, despite different release times. This may indicate a topographically controlled front which forms commonly on the western sides of Heron Reef, noticeable by the long algal blooms of Trichodesmium observed along the reef during daylight hours and coral spawn slicks on the days following spawning events (Fig. S3a). Release of drogues near the site on flood and ebb tides indicated that tidal currents ran parallel to the reef slope (Fig. S3c).

**Fig. 1.**
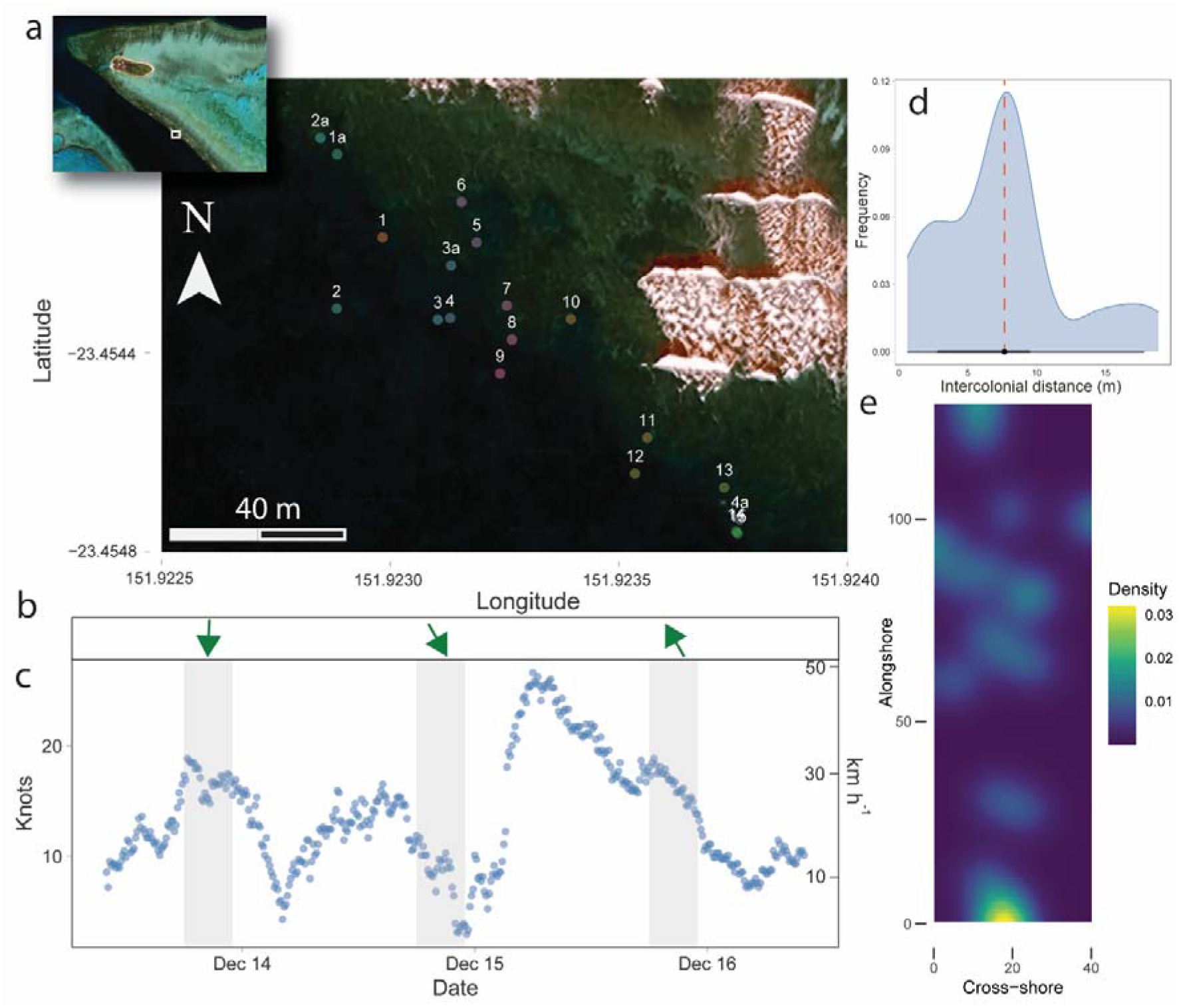
Patch and environmental conditions during the spawning period. (a) Patch distribution of gravid adult Platygyra daedalea colonies along the reef slope (2–9 m depth). (Inset: The western side of Heron Reef and field site.) (b) Wind direction (green arrows) and (c) wind speed during the spawning period. Grey boxes denote nightly spawning periods. (d) Intercolonial distance and spatial distribution densities of gravid colonies within the patch. The red line indicates median intercolonial distance. The low bar indicates the 66% (thick) and 95% (thin) probability intervals. (e) Density of gravid colonies within the patch showing the degree of spatial clustering.

**Table 1.** Meteorological conditions during the nights of the experiment (Barneche et al. 2021).

| Spawning date | Predicted tidal height (m) | Tide direction | Tide amplitude (m) | Wind speed (knots) | Wind direction (°) | Last light |
| --- | --- | --- | --- | --- | --- | --- |
| 13 Dec | 1.2 | Early flood | 1.4 | 14 (17)* | N (6) | 18:58 |
| 14 Dec | 1.3 | Slack low | 1.3 | 7 (9)* | NW (333) | 18:59 |
| 15 Dec | 1.2 | Slack low | 1.3 | 15 (17)* | SSE (165) | 18:59 |

\*Denotes maximum wind speed.

### Patch characteristics

Of the gravid P. daedalea colonies along the transect, the median intercolonial distance was 7.7 m, and the density was 0.004 gravid individuals m^-2^. The Clarke-Evans index was 0.784, indicating spatial clustering (Fig. 1e). After considering genetic groupings (see below), Group 1 resulted in a median intercolonial distance of 14.4 m and a density of 0.002 gravid individuals m^-2^, whereas Group 2 resulted in a median intercolonial distance of 7.99 m and a density of 0.003 gravid individuals m^-2^. Corals within the transect spawned asynchronously across days and time from last light (Table S1). Twelve of the fifteen colonies showed visible evidence of spawning during the monitoring period, and the remaining three colonies were missing eggs upon reassessment.

Although there was a strong positive relationship (r = 0.993) between spawning times of individuals (n=4) and their intercolonial distances on the first spawning night, the relationship was not statistically significant (p = 0.250), likely owing to the low statistical power due to few replicates. The second night of spawning (n=6) showed no significant relationship between spawning times and intercolonial distance (r = 0.112, p = 0.186). When genetic groups were also assessed for spawning times using ‘early’ or ‘late’ binning, all individuals in Group 1 spawned late (i.e. >45 minutes after sunset), whereas five of seven individuals in Group 2 spawned early (p = 0.061). There was no relationship between Group 1 and Group 2 with depth (t = 0.898, p = 0.394).

### Fertilisation success

Due to many corals spawning outside of the diver monitoring periods, combined with the loss of two containers, only five containers were successfully released and collected. The mean fertilisation success in the experimental samples was 1.5%. While self-fertilisation in the field control samples (n =3) was 0%, all individuals that were successfully sequenced (n = 28) were identified as selfed embryos via paternity assignments under both CERVUS and COLONY analyses, closely matching the genetic characteristics of their known egg donor (Table S2-3, Fig. 2). Further, there was a relatively significant decrease in heterozygosity as indicated using FIS between known parents and offspring (t = -6.3858, df = 6, p-value < 0.001).

**Fig. 2.**
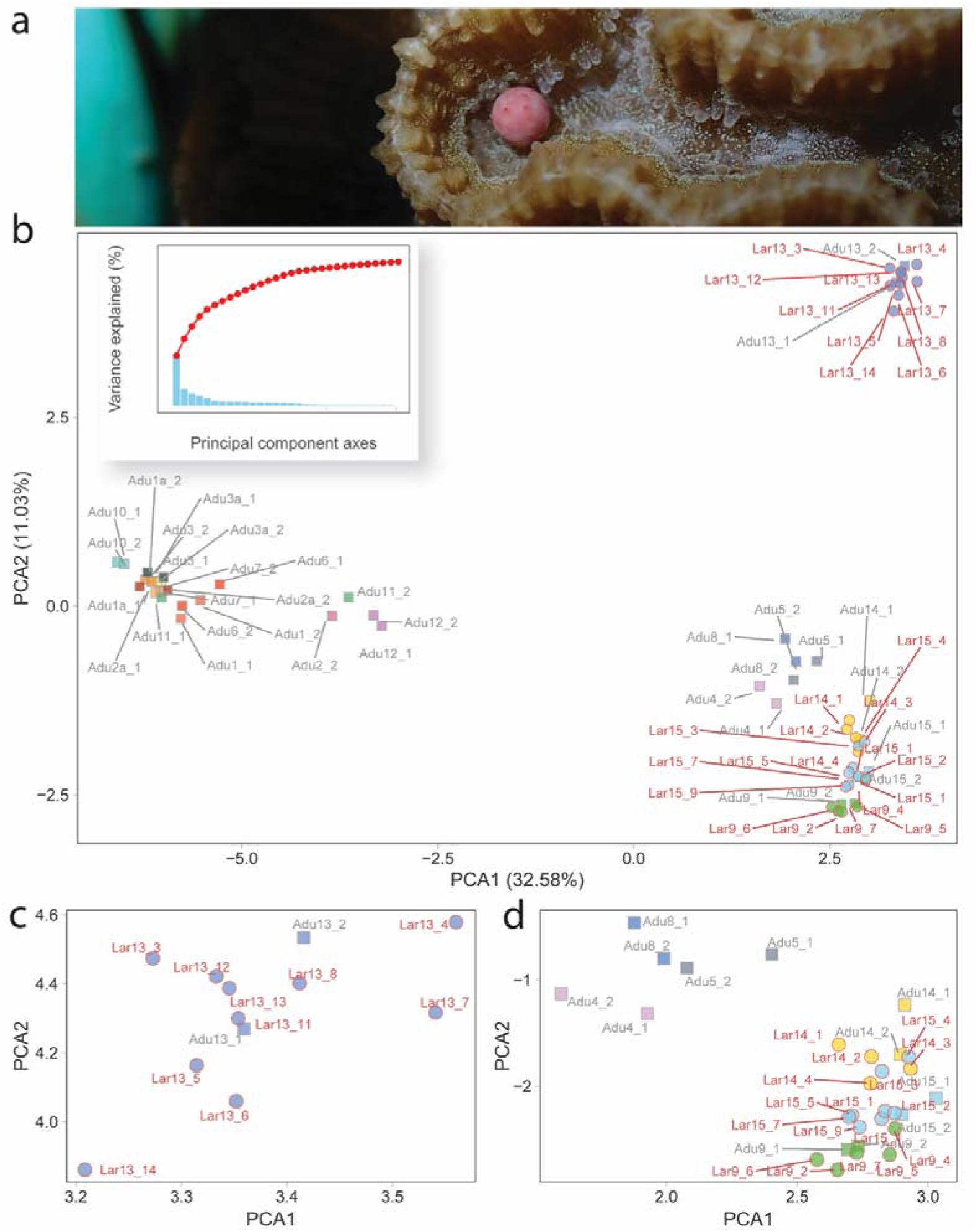
(a) Platygyra daedalea releasing an egg-sperm bundle during spawning. (b) Principal Component Analysis (PCA) of adults and larvae. Square borders denote adults, circle borders denote larvae. Each genotype is coloured individually, and larval colouring matches that of known egg donor colony genotype. Labelling corresponds to colony identity sample number and replicate number. (Inset) Variance explained by each principal component axis. (c) A subset plot of the top-right cluster and (d) a subset plot of the bottom-right cluster.

Further, the laboratory control for selfing where eggs from a single individual were exposed to its own sperm at >10^6^ sperm mL^-1^ for an extended contact time, resulted in low self-fertilisation (<1%) at 3-hours post-spawning. However, at six hours the controls were again assessed and found to be 15% fertilised. All the selfed embryos at this census time point were in the two-cell stage, indicating self-fertilisation was delayed. Contamination of sperm from other individuals was not possible as only one colony spawned ex situ on that night. There was no evidence of parthenogenesis in the eggs washed free of sperm.

### Genetics and colony contribution

Adults were clustered into two main groups (herein Group 1 and Group 2), with colony 12 separated as a ‘noise’ group due to the limited samples (Group 3) (Fig. 3a). STRUCTURE analysis initially indicated the presence of two genetic clusters (K = 2). However, we chose to analyse the structure at K = 3 to account for Group 3, which likely was not identified as a separate group due to the small number of individuals sampled. Colony 13 was not observed in a separate group (Fig. 3a), as seen earlier in the combined adult and larvae clustering (Fig. 2b). Interestingly, colony 12 showed substantial membership from an unrelated group other than genetic Group 1 or 3 (Fig. 3b). There were no clear morphological features that could separate the two main groups by visual inspection (Fig. S4). The inbreeding coefficient (F) values had a median of 0.499, indicating a high probability of inbreeding within the population.

**Fig. 3.**
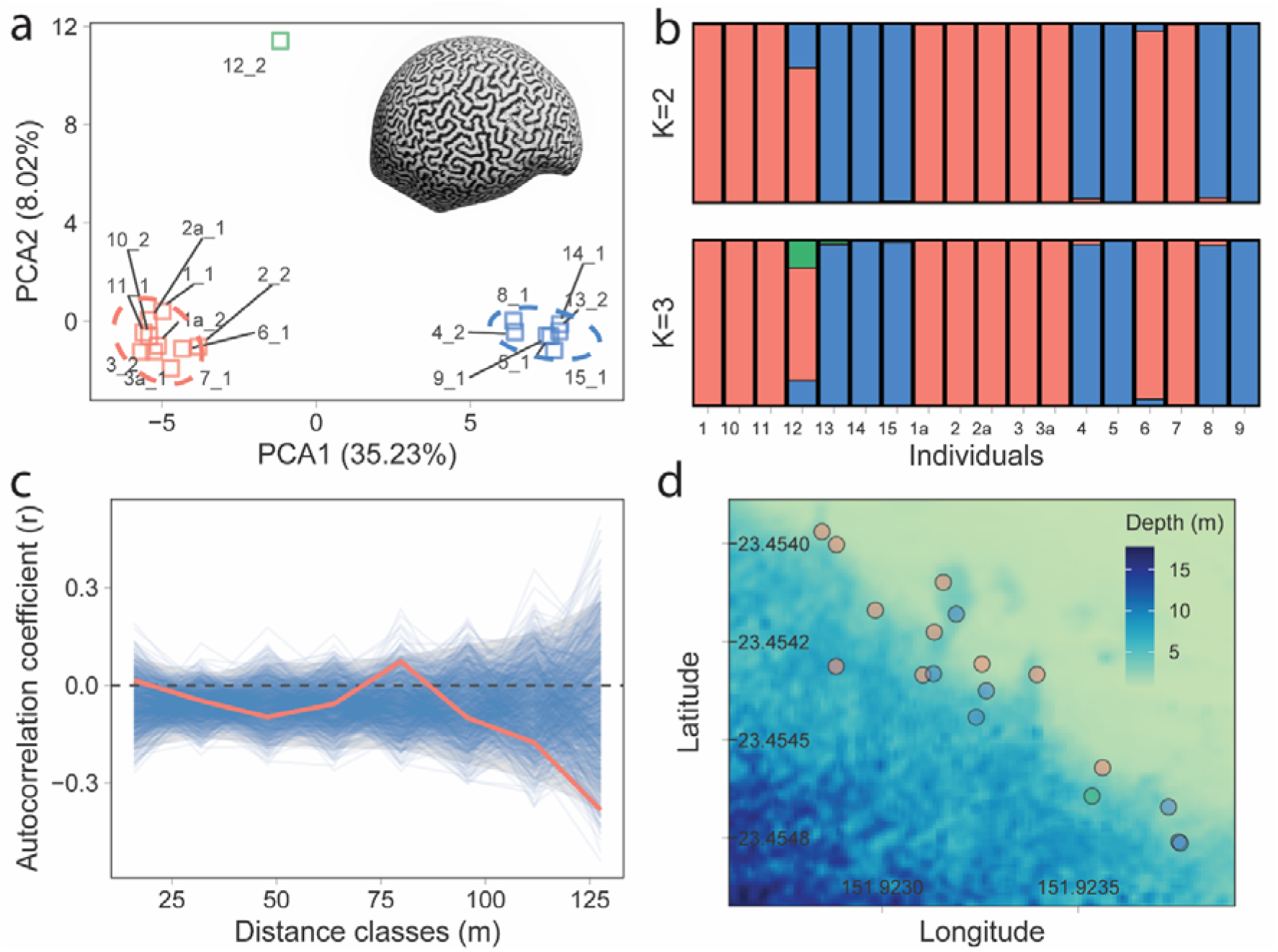
Genetic clustering and spatial relationships among the adult Platygyra daedalea colonies within the transect. (a) Principal Component Analysis (PCA) ordination of unique adult colonies. Group 1 = blue. Group 2 = red. Group 3 = Green. (b) Admixture plots using STRUCTURE for K = 2 and K = 3 clusters. (c) Spatial autocorrelation plot between spatial distance and genetic relatedness. Moran’s I autocorrelation coefficient (r) for different distance classes (m). Red line = raw values. Blue lines = permutations. Grey ribbon = 95% CI of permutations. (d) The adult distribution in genetic clusters relative to depth (bathymetry map from Hedley (2012)).

There was a low, although highly significant, effect of linkage disequilibrium assessed via Index of Association (IA) (*r̄*_*d*_ = 0.047, p = 0.001) suggesting some degree of linkage between the loci in the dataset among the whole group. However, after grouping individuals into subpopulations, there were no significant effects for either group (Group 1: *r̄*_*d*_ = 0.002, p = 0.590; Group 2: 0.011, p = 0.859). Further analysis revealed that each of the 18 adult individuals possess a unique multilocus genotype (MLG), indicating an absence of direct clonality (fragments) in the adult dataset.

Spatial autocorrelation analysis was marginally significant at the furthest distances between individuals within the patch, with a negative r occurring at ∼90 m distances (Fig. 3c). This suggests a pattern of genetic differentiation at larger distances. Similarly, genetic groupings appeared spatially separated along the reef slope, but independent of depth (Fig. 3d).

## Discussion

Despite the wide range of adult colony densities within our patch, outcrossing in P. daedalea was absent along this section of reef. This was driven by a confluence of factors: (i) corals spawning asynchronously across months, characterised by a split spawning event where only approximately one in four colonies were gravid across the reef slope, (ii) asynchronous spawning across many nights, further contributing to a fragmented reproductive pattern, and (iii) asynchrony within the same night. This pronounced asynchrony across multiple temporal scales ultimately implies that a relatively limited number of gametes encountered sufficient sperm concentrations necessary for successful outcrossed fertilisation. This was corroborated by the low fertilisation success observed in our collected samples, with no evidence of outcrossed embryos. While our study did not experimentally manipulate colony density, the observed relationship between asynchrony, low gamete encounters, and reduced fertilisation success is consistent with an Allee effect, where reproductive success declines due to insufficient gamete availability. Recent studies using similar methods, where fertilisation was tracked from contained eggs sourced from natural and manipulated adult patches, have reported up to 30% outcrossing success in synchronous, high-density spawners (Mumby et al. 2024, Ricardo et al. 2024). This comparison reinforces that the low success in our patch was likely driven not simply by adult population density alone but also by the compounding effects of temporal asynchrony on fertilisation.

Unusual spawning patterns within single nights have been documented in Platygyra previously (Miller 1994b), and we observed one colony spawning multiple times within a single night. However, ordination- and model-based clustering analyses of individuals into two main genetic groups reduced the spawning variability within nights, indicating these two groups may be mostly reproductively isolated through distinct spawning timings and possibly undergoing initial stages of allochronic speciation. Here, we avoid using the term ‘cryptic’ species without further population or reproductive data (Patterson et al. 2006, Ramírez-Portilla et al. 2022). However, morphological stasis is known to mask ecological divergence (Bongaerts et al. 2021), and while our population of sampled P. daedalea morphologies all fell within the expected range of morphologies for this species, distinct genetic groupings and spawning patterns are apparent, providing evidence that ecological divergence is occurring.

Although caution is needed when comparing species with complex taxonomy across studies, the observed intercolonial spacing and density of gravid P. daedalea colonies were substantially lower than total adult densities reported in previous studies. Even without considering the increased spacing resulting from two genetic groups, the mean intercolonial distance of gravid colonies was 7.7 m. These spacings are markedly greater than the 1.2 m distances historically reported for this species on the northern Heron reef flat (Endean et al. 1997). Moreover, when considering gravid colonies only, densities were found to be 71-fold lower than historic densities on the northern reef flat, and 8.4 to 19.5 times lower than at nearby One Tree Island (Miller & Ayre 2008). These findings suggest that while split spawning events are theorised to enhance the reliability of larval supply (Hock et al. 2017), they may also contribute to substantial gamete waste (Knowlton 2001). Contrary to the regular spacing of merulinids observed by Endean et al. (1997), we found a tendency towards spatial clustering in gravid P. daedalea colonies. This clustering may be attributed to the spur and groove formations in this section of reef, which likely constrains the availability of suitable habitat space compared to densities reported on the reef flat.

While self-fertilisation in many broadcast spawning corals is typically low, elevated levels of selfing have been documented in Platygyra and other genera (Miller 1994b, Willis et al. 1997). In laboratory settings, the capacity for self-fertilisation may be overestimated due to artificially high sperm concentrations, prolonged egg-sperm contact times, and the absence of competing conspecific sperm. In this study, we found that the few embryos in our samples were genetically close to their maternal parent and exhibited higher homozygous alleles, indicating selfing despite the dilutive conditions and possible sperm competition present in the field. Although our field controls initially detected no evidence of self-fertilisation, laboratory controls—where eggs were exposed to saturating self-sperm concentrations for extended periods—revealed a delayed yet substantial level of fertilisation success. Initial fertilisation was minimal at three hours but significantly increased by 6- hours, coinciding with the onset of primary cell cleavage, usually observed within 90 to 120 minutes following spawning. The two-cell stage observed at six hours suggests fertilisation instead occurred at approximately four- to five-hours post-spawning. While we cannot disregard the likelihood that these embryos resulted from gynogenetic parthenogenesis, which allows some level of genetic recombination, this phenomenon has not been described in this species and we did not observe parthenogenesis in our laboratory experiments, therefore, we consider it unlikely. The absence of self-fertilisation in field controls likely reflects that these samples were counted early, prior to the commencement of delayed selfing or that not all individuals have the capacity to self.

Delayed self-fertilisation has been documented in several coral species in environments where conspecific sperm are low or absent (Heyward & Babcock 1986, Willis et al. 1997, Isomura et al. 2016) and combines the advantages of outcrossing and selfing i.e., a bet-hedging strategy to produce progeny if preferred strategy fails (Goodwillie & Weber 2018). The mechanism for delayed selfing is poorly studied in marine invertebrates compared with higher plants (Kalisz et al. 1999, Rea & Nasrallah 2008), but results from the relaxation of the self-incompatibility (SI) system in the gametes as they age (Goodwillie & Weber 2018). In ascidians, gene expression in the egg’s vitelline coat, as well as the use of self-incompatibility molecules have been proposed as a mechanism for self/nonself-recognition (Saito & Sawada 2022). Additionally, we observed high inbreeding coefficients in the adult population, which has been linked with an organism’s ability to self (Olsen et al. 2020), and may indicate that selfing plays an influential role in shaping the genetic structure of this population.

Contrary to the findings of Miller and Ayre (2008), we did not observe clonal individuals within our transect, even among those in close proximity (e.g. Adults 14 and 15). However, inbreeding coefficients were high and inbreeding has been suggested to be common in marine invertebrates, even those that disperse gametes and randomly mate (Olsen et al. 2020). The spatial autocorrelation analysis revealed a negative, marginally significant, trend between intercolonial distance and genetic relatedness across a fine-spatial scale. Interestingly, this trend across fine scales has been observed before, even when clones are accounted for (Miller & Ayre 2008, Dubé et al. 2020). Here, negative relatedness values become more pronounced at intercolonial distances of approximately 90 m. This pattern, typical of brooding species whose larvae often settle near the parent colony (Prata et al. 2024), was highly unexpected in broadcast spawners where localised stock-recruitment relationships are less established in open populations (Doropoulos et al. 2015), but see Ayre and Hughes (2000), Japaud et al. (2019), Dubé et al. (2020). However, mobile bodies resembling larvae have been visually observed within polyps of a Platygyra sp. (R. Babcock pers. comm.), indicating some species in this genus may be undescribed facultative brooders.

An alternative hypothesis to explain this spatial-genetic correlation is that larvae may preferentially settle near conspecific adult groups. While this hypothesis contrasts with some existing observations of Janzen-Connell effects, a theory that has been extensively studied in terrestrial systems, these effects have only recently been explored in marine invertebrates at small spatial scales of metres and may require further examination at scales of tens to hundreds of metres (Johnson et al. 2012, Marhaver et al. 2013, Sims et al. 2021). Additionally, the possibility of larvae settling as sibling groups through collective dispersal, although less plausible, could explain the observed spatial patterns (see Eldon et al. (2016) for a review). This scenario depends on cross-fertilisation among nearby spawning adults (Levitan et al. 2004, Ricardo et al. 2024), followed by the settlement of siblings in close proximity to one another. Despite short larval competency times of P. daedalea (Miller & Mundy 2003) and that drifter releases close to the study site indicated that larvae may be retained locally, and possibly within the Heron-Wistari channel, divergence of mesh container trajectories even within a few hours of spawning indicates this hypothesis is unlikely. Few studies have assessed siblingship patterns in corals, and further work is needed to assess if such patterns are common or play a meaningful role in shaping spatial-genetic structures (Puill-Stephan et al. 2009, Dubé et al. 2020).

Platygyra lacks distinct morphological and reproductive traits that allow clear delineation into species i.e., a syngameon. Although the majority of the P. daedalea colonies were identified as the Classic PDC P. daedalea morphotype (Miller 1994a), clear morphological differences were observed even within this morphotype. Nonetheless, no consistent morphological correlation with our genetic partitioning of two main groups was observed. Colony 12 was a distinct morphological variant (‘fat morph’; K. Miller pers. comm.), but there was only one individual within our transect and therefore we could not compare it against other groups. While some morphotypes within P. daedalea have shown reduced cross-fertilisation success (Miller 1994b), this is unlikely to be the cause of the absence of outcrossing in our experiment because almost all corals in our transect appeared to be the same morph. The genetic clustering we observed contrasted that observed in Miller and Benzie (1997), who observed little genetic differentiation in the genus Platygyra. The contrast in findings may be due to true differences in fine-scale population structure between the studies, but may also be a result of methodological differences i.e., SNP genotyping used here typically resolves greater levels of grouping than observed in microsatellites (Zimmerman et al. 2020, Pérez-González et al. 2023).

It is clear that low sexual outcrossing is common in Platygyra during split spawning events. Reproductive plasticity allowing self-fertilisation may provide a minor advantage by producing a limited number of offspring instead of wasting all gametes when sexual outcrossing fails during low-density spawning events. Although clones were not observed in the samples or transect, as has been observed elsewhere (Babcock 1991, Miller & Ayre 2008), there was some evidence of inbreeding or asexual reproduction in the adult population data. Together, these findings suggest that, without an unidentified reproductive mechanism or hydrodynamic convergence zones, split spawning events on typical reef tracts may lead to substantial gamete wastage – a common feature even in high density slicks (Hodgson 1988, Simpson et al. 1993). The contribution of outcrossed larvae may be largely restricted to areas of higher colony densities, both at the reef-scale and across the metapopulation, where certain dense populations contribute disproportionately, indicating that effective spawning densities must surpass the low levels observed here. Restoration efforts aimed at enhancing functional reproduction on reefs should consider adult densities but also reproductive modes, spawning synchrony, and inherent population structure to maximise their effectiveness. Moreover, density-dependent processes on reefs, starting with the reproduction of mass spawners that rely on external fertilisation, need to be considered in model simulations of reef dynamics. It is highly likely that most simulation modelling efforts to date have not considered such processes and have thus overestimated larval production during the lag phases of recovery.

## Data availability statement

The datasets used for this study are deposited in the CSIRO Data Access Portal https://data.csiro.au/ and will be made fully available following publication.

## Acknowledgments

The authors would like to acknowledge the Traditional Owners of the Great Barrier Reef, particularly the Byelle, Gooreng Gooreng, Gurang and Taribelang Bunda First Nations people of the Port Curtis Coral Coast, and the Manbarra First Nations people of the Palm Islands, for permission to work in their Sea Country with free prior and informed consent. We pay our respects to their Elders, past, present, and emerging, and acknowledge their continuing spiritual connection to their Sea Country. All work was conducted under Great Barrier Reef Marine Park Authority permits G21/44774.1 and G22/46963.1. We thank R. Mason, M. Tonks, and the staff of Heron Island Research Station for field and coordination support, including A. Donovan. We thank M. Miller, A. Lamb, and I. Popovic for discussions on genetics, and F. Sjahruddin for providing the wave data. This work was supported by the EcoRRAP subprogram (https://gbrrestoration.org/program/ecorrap/) that is part of the Reef Restoration and Adaptation Program (RRAP, https://gbrrestoration.org/). RRAP is funded by the partnership between the Australian Governments Reef Trust and the Great Barrier Reef Foundation. The funders had no role in the study design, data collection and analysis, decision to publish, or preparation of the manuscript.

## CRediT author statement

Conceptualization: Peter Mumby, Christopher Doropoulos, Gerard Ricardo Methodology: Gerard Ricardo, Peter Mumby, Russell Babcock, Christopher Doropoulos Formal analysis: Gerard Ricardo, Arne Adam

Investigation: Gerard Ricardo, Peter Mumby, Russell Babcock, Natalia Robledo, Elizabeth Buccheri, Julian Uribe Palomino, Christopher Doropoulos

Writing - Original Draft: Gerard Ricardo Writing - Review & Editing: All authors Visualization: Gerard Ricardo

Supervision: Peter Mumby, Christopher Doropoulos Funding acquisition: Christopher Doropoulos, Peter Mumby

## Notes

### Competing Interest Statement

The authors have declared no competing interest.

### Summary of Updates

Some changes in terminology were made.

