## Supplementary Material for "Spawning asynchrony and mixed reproductive strategies in a common mass spawning coral"

### Supplementary materials

Table S1. Individual times of spawning of *Platygyra* *daedalea* across spawning nights. DAFM = Days After the Full Moon. Brackets denote the number of colonies spawning at the timepoint. Times are standardised to last light (6:58 pm). Note: some colonies spawned on multiple days and twice on the same night.

| Morphotype | 5 DAFM | 6 DAFM | 7 DAFM |
| --- | --- | --- | --- |
| PDC (Classic) | 0:52 (3), >2:52 (1) | -0:12 (2), -0:06 (2), 0:50 (1) | >0:38 (4) |
| PH (Hillocky) | – | -0:16 (1) | – |
| PDSE (Encrusting) | – | – | >0:38 (1) |

Table. S2. CERVUS output for paternity assignments. Two replicate samples of candidate sperm donor colonies were provided in the analysis.

| Offspring ID | Egg donor ID | Candidate sperm donor ID | Trio LOD score | Trio confidence |
| --- | --- | --- | --- | --- |
| pd13_l_14_11 | pd13_a_1 | pd13_a_2 | 9.08E+01 | * |
| pd13_l_14_12 | pd13_a_1 | pd13_a_2 | 8.69E+01 | * |
| pd13_l_14_13 | pd13_a_1 | pd13_a_2 | 1.03E+02 | * |
| pd13_l_14_14 | pd13_a_1 | pd13_a_2 | 7.41E+01 | * |
| pd13_l_14_16 | pd13_a_1 | pd13_a_1 | 2.25E+00 | * |
| pd13_l_14_3 | pd13_a_1 | pd13_a_2 | 1.04E+02 | * |
| pd13_l_14_4 | pd13_a_1 | pd13_a_2 | 1.06E+02 | * |
| pd13_l_14_5 | pd13_a_1 | pd13_a_2 | 1.02E+02 | * |
| pd13_l_14_6 | pd13_a_1 | pd13_a_2 | 8.47E+01 | * |
| pd13_l_14_7 | pd13_a_1 | pd13_a_2 | 1.01E+02 | * |
| pd13_l_14_8 | pd13_a_1 | pd13_a_2 | 1.01E+02 | * |
| pd14_l_14_1 | pd14_a_1 | pd14_a_1 | 1.12E+02 | * |
| pd14_l_14_2 | pd14_a_1 | pd14_a_1 | 1.23E+02 | * |
| pd14_l_14_3 | pd14_a_1 | pd14_a_1 | 1.22E+02 | * |
| pd14_l_14_4 | pd14_a_1 | pd14_a_1 | 1.32E+02 | * |
| pd15_l_13_1 | pd15_a_1 | pd15_a_2 | 1.05E+02 | * |
| pd15_l_14_1 | pd15_a_1 | pd15_a_2 | 8.94E+01 | * |
| pd15_l_14_2 | pd15_a_1 | pd15_a_1 | 9.34E+01 | * |
| pd15_l_14_3 | pd15_a_1 | pd15_a_2 | 9.84E+01 | * |
| pd15_l_14_4 | pd15_a_1 | pd15_a_2 | 9.97E+01 | * |
| pd15_l_14_5 | pd15_a_1 | pd15_a_1 | 1.00E+02 | * |
| pd15_l_14_7 | pd15_a_1 | pd15_a_2 | 9.59E+01 | * |
| pd15_l_14_9 | pd15_a_1 | pd15_a_1 | 8.85E+01 | * |
| pd9_l_14_2 | pd9_a_1 | pd9_a_1 | 1.23E+02 | * |
| pd9_l_14_4 | pd9_a_1 | pd9_a_1 | 1.30E+02 | * |
| pd9_l_14_5 | pd9_a_1 | pd9_a_2 | 1.19E+02 | * |
| pd9_l_14_6 | pd9_a_1 | pd9_a_1 | 1.12E+02 | * |
| pd9_l_14_7 | pd9_a_1 | pd9_a_1 | 1.17E+02 | * |

Table. S3. COLONY output for paternity assignments Two replicate samples of candidate sperm donor colonies were provided in the analysis.

| OffspringID | Inferred_Sperm_donor | Inferred_Egg_donor | Probability |
| --- | --- | --- | --- |
| pd13_l_14_3 | pd13_a_1 | pd13_a_1 | 0.9999 |
| pd13_l_14_4 | pd13_a_1 | pd13_a_1 | 1 |
| pd13_l_14_5 | pd13_a_1 | pd13_a_1 | 1 |
| pd13_l_14_6 | pd13_a_1 | pd13_a_1 | 0.9999 |
| pd13_l_14_7 | pd13_a_1 | pd13_a_1 | 0.9999 |
| pd13_l_14_8 | pd13_a_1 | pd13_a_1 | 0.9999 |
| pd13_l_14_11 | pd13_a_1 | pd13_a_1 | 1 |
| pd13_l_14_12 | pd13_a_1 | pd13_a_1 | 1 |
| pd13_l_14_13 | pd13_a_1 | pd13_a_1 | 1 |
| pd13_l_14_14 | pd13_a_1 | pd13_a_1 | 0.9999 |
| pd14_l_14_1 | pd14_a_1 | pd14_a_1 | 1 |
| pd14_l_14_2 | pd14_a_1 | pd14_a_1 | 0.9969 |
| pd14_l_14_3 | pd14_a_1 | pd14_a_1 | 0.9991 |
| pd14_l_14_4 | pd14_a_1 | pd14_a_1 | 1 |
| pd15_l_13_1 | pd15_a_1 | pd15_a_1 | 0.9135 |
| pd15_l_14_1 | pd15_a_1 | pd15_a_1 | 0.9143 |
| pd15_l_14_2 | pd15_a_1 | pd15_a_1 | 0.9075 |
| pd15_l_14_3 | pd15_a_1 | pd15_a_1 | 0.9117 |
| pd15_l_14_4 | pd15_a_1 | pd15_a_1 | 0.9124 |
| pd15_l_14_5 | pd15_a_1 | pd15_a_1 | 0.9128 |
| pd15_l_14_7 | pd15_a_1 | pd15_a_1 | 0.9139 |
| pd15_l_14_9 | pd15_a_1 | pd15_a_1 | 0.9134 |
| pd9_l_14_2 | pd9_a_1 | pd9_a_1 | 0.7497 |
| pd9_l_14_4 | pd9_a_1 | pd9_a_1 | 0.7944 |
| pd9_l_14_5 | pd9_a_1 | pd9_a_1 | 0.7998 |
| pd9_l_14_6 | pd9_a_1 | pd9_a_1 | 0.5071 |
| pd9_l_14_7 | pd9_a_1 | pd9_a_1 | 0.9683 |

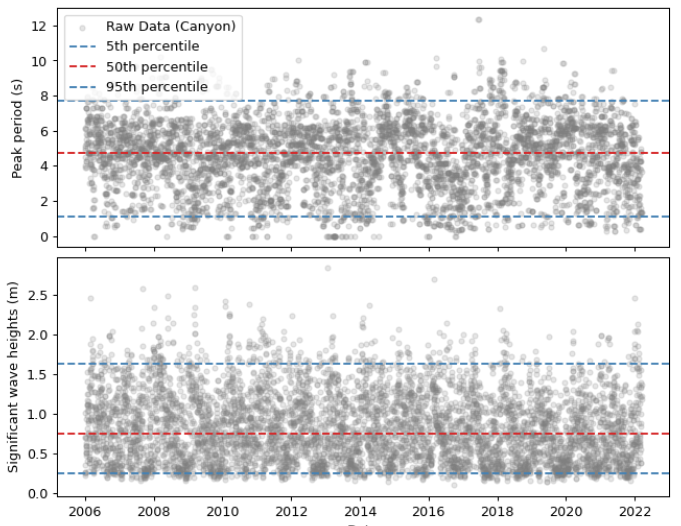

Fig. S1. Modelled wave time series data from 2005–2022 at the field site (Canyons, Heron Island) including peak period and significant wave heights (Callaghan 2023).

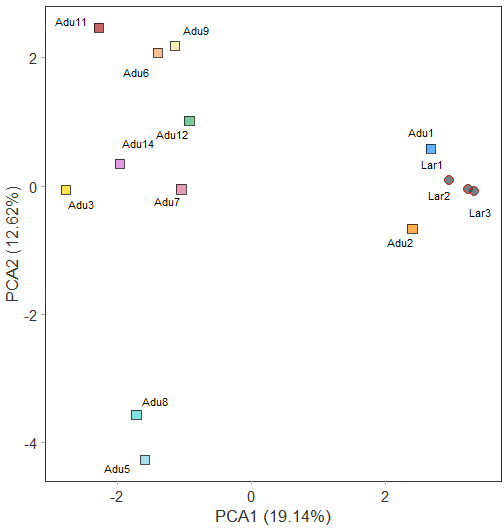

Fig. S2. Principal Components Analysis (PCA) of adult candidate parents (n = 11) and larvae (n = 3) of *A. kenti*. Larvae 1 to 3 belonged to known parents, Adults 1 and 2.

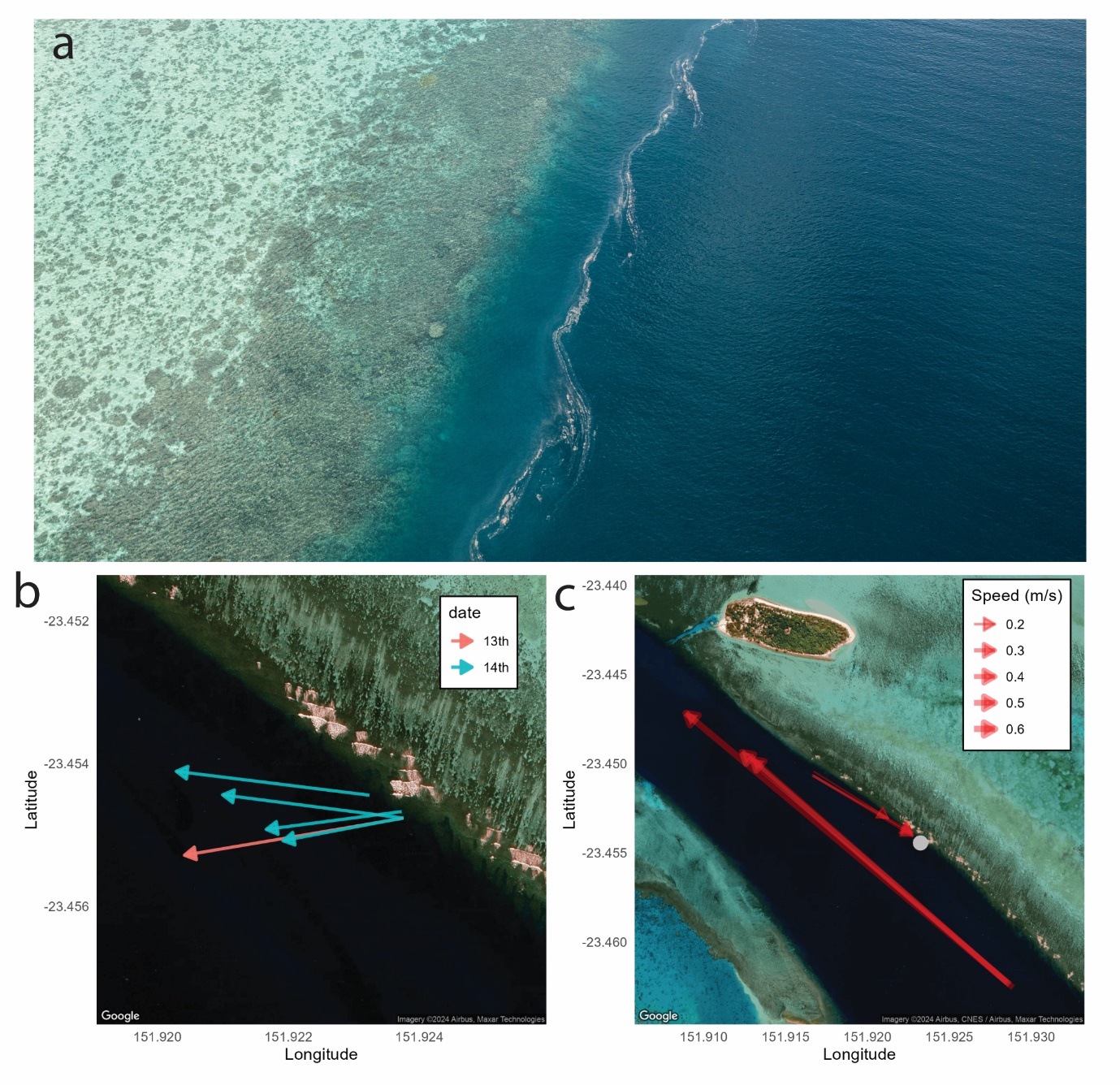

Fig. S3. Hydrodynamics of the field site. (a) A topographically controlled front indicated by a coral spawn slick on the western side of Heron reef in 2021 on the day following spawning (© CSIRO. Credit: Nick Thake 2021). (b) Movement of the mesh containers following release for each spawning night. (c) Movement and speed of drogues released within the Heron-Wistari channel. The grey point denotes the field site.

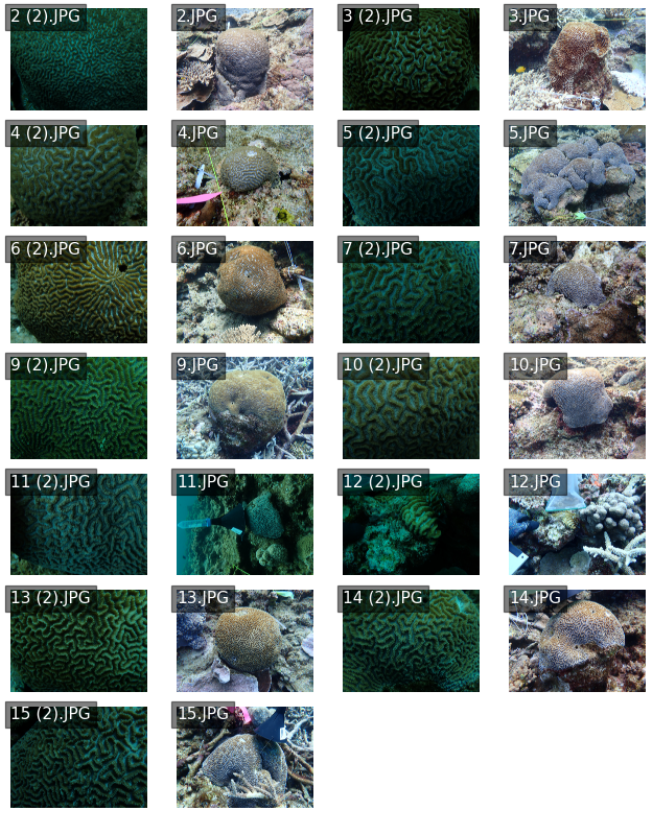

Fig. S4. Images of adult colonies of *Platygyra daedalea* tagged within the site. Higher resolution images can be accessed at <https://doi.org/10.6084/m9.figshare.27934917>.
